# Novel Computational Models of Evoked Dopamine Release In Vivo Measured by Fast Scan Cyclic Voltammetry Quantify the Regulation of Presynaptic Kinetics by Synucleins

**DOI:** 10.1101/2022.05.04.490695

**Authors:** N Shashaank, Mahalakshmi Somayaji, Mattia Miotto, Eugene V. Mosharov, David A. Knowles, Giancarlo Ruocco, David L. Sulzer

## Abstract

Dopamine neurotransmission in the striatum is central to many normal and disease functions. Ventral midbrain dopamine neurons exhibit ongoing tonic firing that produce low extrasynaptic levels of dopamine below the detection of extrasynaptic electrochemical recordings (∼10 – 20 nanomolar), with superimposed bursts that can saturate the dopamine uptake transporter and produce transient micromolar concentrations. The bursts have previously been shown to lead to presynaptic plasticity via multiple mechanisms, but analysis methods for these kinetic parameters are limited. To provide a deeper understanding of the mechanics of dopamine neurotransmission, we present three computational models of dopamine release with different levels of spatiotemporal complexity to analyze in vivo fast-scan cyclic voltammetry recordings from the dorsal striatum of mice. The models accurately fit to the cyclic voltammetry data and provide estimates of presynaptic dopamine facilitation/depression kinetics and dopamine transporter reuptake kinetics. We use the models to analyze the role of synuclein proteins in neurotransmission and quantify recent findings linking presynaptic protein α-synuclein to the short-term facilitation and long-term depression of dopamine release.

## Introduction

Dopamine (DA) is a neurotransmitter that plays important roles in learning and memory, drug and alcohol addiction, attention deficit disorder, schizophrenia, and neurodegenerative disorders including Parkinson’s disease^1^. Electrochemical recording methods, particularly fast-scan cyclic voltammetry (FSCV) and constant potential amperometry, are used to record and analyze evoked DA release from the axons of dopamine neurons in rodent brains in vivo^2,3^, ex vivo in brain slice preparations^4,5^, and in cultured neurons^6,7^. The development of computational models to quantitatively describe FSCV and amperometry recordings of evoked DA release^8–12^ has enabled the detailed analysis of DA release and reuptake kinetics under different stimulation paradigms both ex vivo and in vivo. However, the plasticity of DA neurotransmission is far more complex in vivo due to multiple factors that are less important in culture or slice preparations, including modulatory inputs from neurotransmitters such as acetylcholine and γ-aminobutyric acid (GABA) and multiple presynaptic regulatory proteins such as the synapsins and synucleins. Importantly, ventral midbrain DA neurons are tonic pacemakers, but their cell bodies are absent in slice preparations.

In vivo, tonic firing produces minimal DA levels in the striatum and is interspersed with burst firing activity that can cause DA release to overwhelm the dopamine transporter (DAT) reuptake system, thus providing supralinear levels of extracellular DA during bursts. As reported in a study of spontaneous DA release^9^, the magnitude and duration of burst firing-evoked DA release in vivo are subject to multiple depressing and facilitatory influences that remain mostly uncharacterized. As such, deciphering the kinetics of DA release during burst firing is central for elucidating the interactions of DA in vivo and providing insights into the mechanisms responsible for DA release and regulation in normal development and function, exposure to drugs (ranging from antipsychotics to recreational/abused drugs), and many disease states.

Developing a general computational model of DA release requires accounting for multiple biological, electrochemical, and experimental factors that affect the FSCV trace. We have created three mathematical models of burst-firing striatal DA release in vivo, each of which closely fits the data and can be used to derive biologically interpretable parameters such as estimates of facilitation/depression kinetics, amount of DA release, and DAT reuptake kinetics. These models can be adapted for other approaches to detect extracellular DA^13^ and for analyzing release kinetics of other neurotransmitters.

The models simulate the interactions between extracellular DA released into striatal tissue, the carbon-fiber electrode that detects DA, and an intermediate area of damaged tissue known as the “dead space,” formed when inserting the electrode into the tissue^14^ (Figure 1A). When an electrical stimulus is applied, DA released by vesicular exocytosis diffuses through the extracellular space towards the dead space and carbon-fiber electrode. In contrast to amperometry which oxidizes DA and acts as a “sink,” FSCV does not consume DA molecules and causes a “bounce back” of DA into the tissue through the dead space^5^. As a result, the concentration of extracellular DA in the striatum with FSCV experiments is mostly regulated by DAT uptake. The concentration of DA measured at the electrode is further altered by electrochemical adsorption^15^, which causes the signal baseline to increase when multiple bursts of stimuli are triggered in quick succession.

**FIGURE 1:**
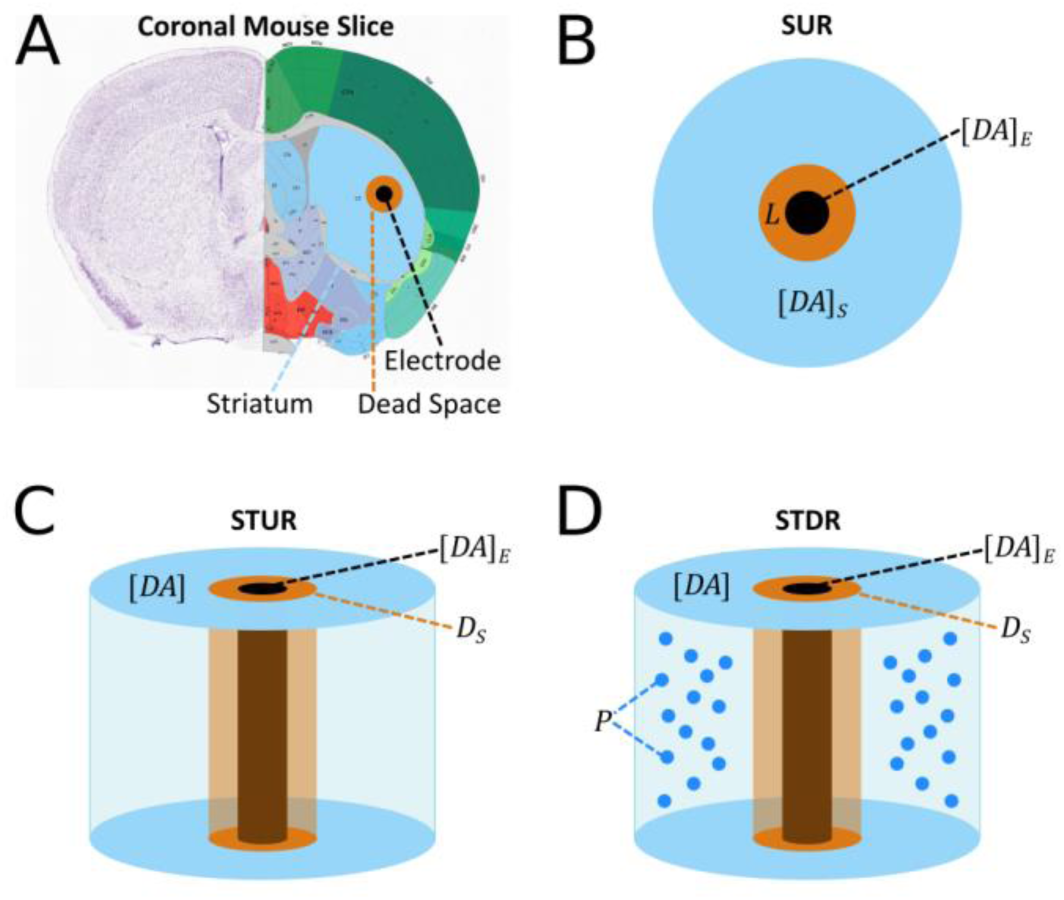
Schematic Representations of Computational Models. **A**, Coronal slice of the mouse brain (adopted from the Allen Brain Atlas). During in vivo FSCV experiments, a bipolar electrode in the ventral midbrain (not shown) is used to electrically stimulate the cell bodies of dopaminergic neurons, causing DA release via vesicular exocytosis from dopaminergic axons in the dorsal striatum (blue). The DA molecules diffuse to the carbon-fiber electrode (black) through striatal tissue and an area of damaged tissue known as the “dead space” (orange) that is formed when the electrode is inserted into the brain. **B**, Simple Uniform Release Model. DA release sites are modeled as a continuous release source within the striatum using temporal diffusion in one dimension without calculating the spatial characteristics of the striatum. **C**, Spatiotemporal Uniform Release Model, which is similar to the Simple Uniform Release model, except that the spatial characteristics of the dorsal striatum are modeled as a cylinder using isotropic diffusion in two dimensions. **D**, Spatiotemporal Discrete Release Model, which is similar to the Spatiotemporal Uniform Release model, with the addition of release sites modeled as discrete point sources positioned throughout the striatum.

While the three models described here differ in the implementation of release and diffusion mechanics, their functional predictions are very similar. The *Simple Uniform Release* (SUR) model (Figure 1B) is 1-dimensional and computes DA release as a uniform distribution in the striatum that diffuses towards the carbon-fiber electrode without taking into account physical characteristics of the striatum. The *Spatiotemporal Uniform Release* (STUR) model (Figure 1C) is 2-dimensional and incorporates spatiotemporal diffusion of DA in the striatum using cylindrical coordinates and a uniform release distribution similar to the SUR model. The *Spatiotemporal Discrete Release* (STDR) model (Figure 1D) is similar to the STUR model but uses discrete DA release sites positioned throughout the striatum. The mathematical equations for each model are described in the Methods section.

## Results

We fit the computational models to in vivo recordings of dorsal striatum DA release evoked by multiple burst stimuli in the ventral midbrain^16^. The complete list of model parameter estimates is provided in the Supplementary Information. Briefly, the experimental procedure was comprised of 6 “Single Burst” and “Repeated Burst” stimulation protocols (each pass being termed a “sweep”), with a 2-minute recovery time between protocols and a 6-minute recovery time between sweeps (Figure 2). The Single Burst protocol was a train of 30 pulses at 50 Hz, and the Repeated Burst protocol was a series of 6 single bursts with a 5 second interstimulus period between two bursts.

**FIGURE 2:**
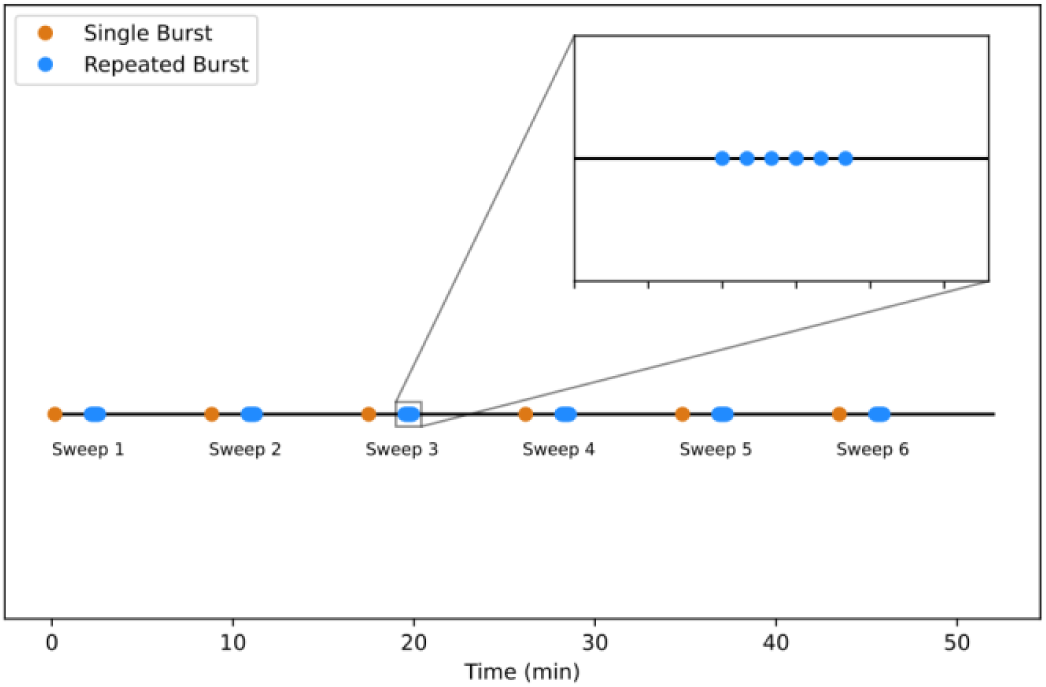
Fast-Scan Cyclic Voltammetry Experiment Stimulus Protocol. The experimental protocol is comprised of 6 sweeps, and each sweep contains a “Single Burst” and “Repeated Burst” protocol, with a 2-minute recovery time between protocols and a 6-minute recovery time between sweeps. A Single Burst (orange) is comprised of 30 pulses at 50 Hz, while a Repeated Burst (blue) is a series of 6 Single Bursts with 5 seconds between each burst.

### Three Kinetic Components Are Sufficient To Describe DA Release Kinetics In Wildtype Mice In Vivo

To analyze the kinetics of DA release under normal conditions, the models were fit to FSCV data from wildtype (WT) mice using the Single Burst and Repeated Burst protocols from Sweep 1 (Figure 3) and Sweep 6 (Figure 4) of the experimental procedure. Consistent with conclusions from a study of spontaneous DA release kinetics in behaving mice^9^, the complex dynamics of the data require three kinetic components to capture changes in DA release over time: short-term facilitation, short-term depression, and long-term depression. Each kinetic is described in the models by a “kick component” *K*_*j*_ that controls the facilitative/depressive effect’s magnitude and a time constant *τ*_*j*_ that controls the kinetic’s duration (see Methods for more information). The kinetic parameter estimates (Table 1) for the Single Burst and Repeated Burst protocols indicate that in WT mice, the kick component for short-term facilitation is 3.5-fold larger than short-term depression, while the time constant of short-term depression is ∼1.7-fold to 2-fold longer than short-term facilitation.

**TABLE 1:**
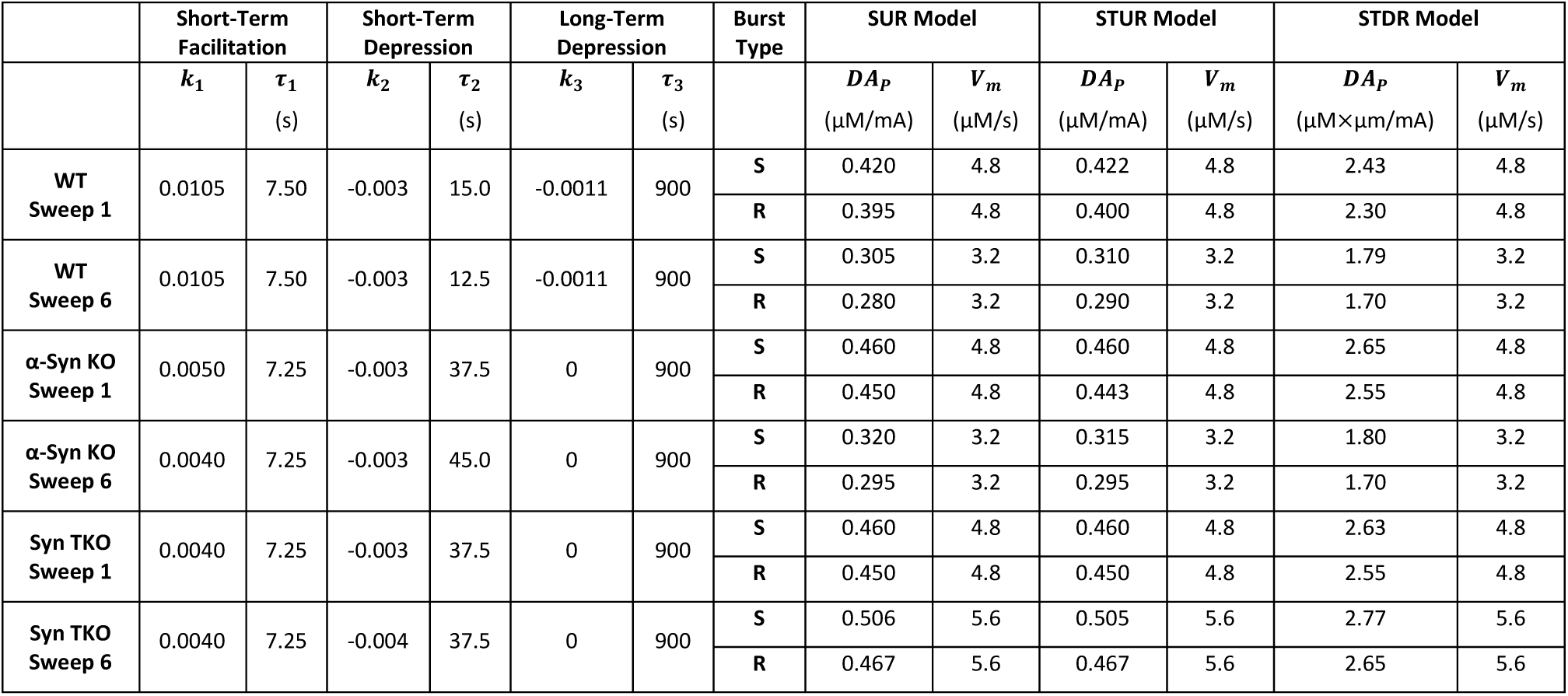
Best Fit Model Parameters for DA Release Kinetics and DAT Activity. For all fits presented in the Results section, *K*_*m*_ was set to 0.2 μM.

|  | Short-Term Facilitation |  | Short-Term Depression |  | Long-Term Depression |  | Burst Type | SUR Model |  | STUR Model |  | STDR Model |  |
| --- | --- | --- | --- | --- | --- | --- | --- | --- | --- | --- | --- | --- | --- |
| | $k_1$ | $\tau_1$<br>(s) | $k_2$ | $\tau_2$<br>(s) | $k_3$ | $\tau_3$<br>(s) | | $DA_p$<br>($\mu\text{M}/\text{mA}$) | $V_m$<br>($\mu\text{M}/\text{s}$) | $DA_p$<br>($\mu\text{M}/\text{mA}$) | $V_m$<br>($\mu\text{M}/\text{s}$) | $DA_p$<br>($\mu\text{M}\times\mu\text{m}/\text{mA}$) | $V_m$<br>($\mu\text{M}/\text{s}$) |
| WT Sweep 1 | 0.0105 | 7.50 | -0.003 | 15.0 | -0.0011 | 900 | S | 0.420 | 4.8 | 0.422 | 4.8 | 2.43 | 4.8 |
|  |  |  |  |  |  |  | R | 0.395 | 4.8 | 0.400 | 4.8 | 2.30 | 4.8 |
| WT Sweep 6 | 0.0105 | 7.50 | -0.003 | 12.5 | -0.0011 | 900 | S | 0.305 | 3.2 | 0.310 | 3.2 | 1.79 | 3.2 |
|  |  |  |  |  |  |  | R | 0.280 | 3.2 | 0.290 | 3.2 | 1.70 | 3.2 |
| α-Syn KO Sweep 1 | 0.0050 | 7.25 | -0.003 | 37.5 | 0 | 900 | S | 0.460 | 4.8 | 0.460 | 4.8 | 2.65 | 4.8 |
|  |  |  |  |  |  |  | R | 0.450 | 4.8 | 0.443 | 4.8 | 2.55 | 4.8 |
| α-Syn KO Sweep 6 | 0.0040 | 7.25 | -0.003 | 45.0 | 0 | 900 | S | 0.320 | 3.2 | 0.315 | 3.2 | 1.80 | 3.2 |
|  |  |  |  |  |  |  | R | 0.295 | 3.2 | 0.295 | 3.2 | 1.70 | 3.2 |
| Syn TKO Sweep 1 | 0.0040 | 7.25 | -0.003 | 37.5 | 0 | 900 | S | 0.460 | 4.8 | 0.460 | 4.8 | 2.63 | 4.8 |
|  |  |  |  |  |  |  | R | 0.450 | 4.8 | 0.450 | 4.8 | 2.55 | 4.8 |
| Syn TKO Sweep 6 | 0.0040 | 7.25 | -0.004 | 37.5 | 0 | 900 | S | 0.506 | 5.6 | 0.505 | 5.6 | 2.77 | 5.6 |
|  |  |  |  |  |  |  | R | 0.467 | 5.6 | 0.467 | 5.6 | 2.65 | 5.6 |

**FIGURE 3:**
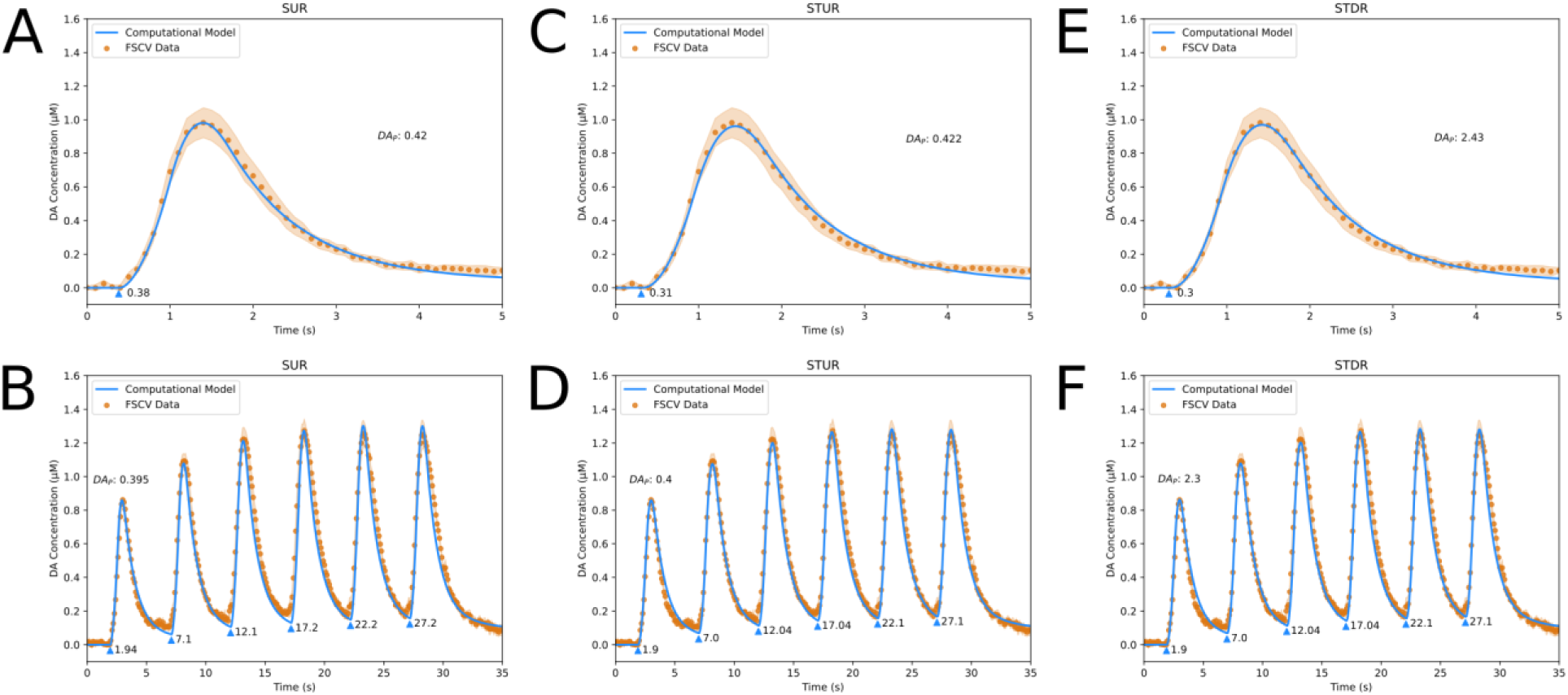
Fits of DA Computational Models to Sweep 1 FSCV Traces from WT Mice in Dorsal Striatum. **A**, fit of Simple Uniform Release Model (*R*^2^ = 0.99) to Single Burst protocol. **B**, fit of Simple Uniform Release Model (*R*^2^ = 0.96) to Repeated Burst protocol. **C**, fit of Spatiotemporal Uniform Release Model (*R*^2^ = 0.99) to Single Burst protocol. **D**, fit of Spatiotemporal Uniform Release Model (*R*^2^ = 0.98) to Repeated Burst protocol. **E**, fit of Spatiotemporal Discrete Release Model (*R*^2^ = 0.99) to Single Burst protocol. **F**, fit of Spatiotemporal Discrete Release Model (*R*^2^ = 0.98) to Repeated Burst protocol. Blue lines are model fits, blue triangles indicate start times *t*_*i*_ of electrical bursts, orange dots are averaged FSCV traces (*N* = 7 animals), and orange ribbons report SEM.

**FIGURE 4:**
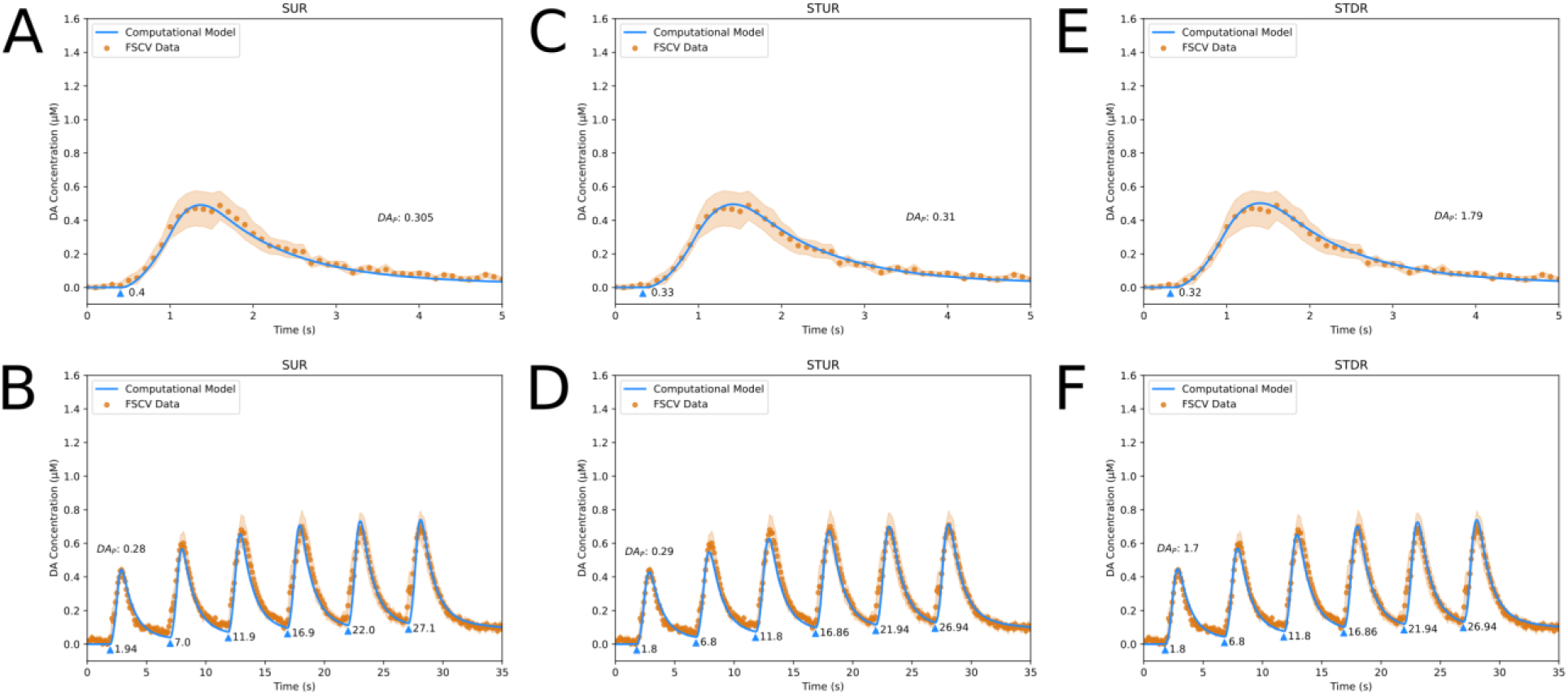
Fits of DA Computational Models to Sweep 6 FSCV Traces from WT Mice in Dorsal Striatum. **A**, fit of Simple Uniform Release Model (*R*^2^ = 0.98) to Single Burst protocol. **B**, fit of Simple Uniform Release Model (*R*^2^ = 0.93) to Repeated Burst protocol. **C**, fit of Spatiotemporal Uniform Release Model (*R*^2^ = 0.99) to Single Burst protocol. **D**, fit of Spatiotemporal Uniform Release Model (*R*^2^ = 0.95) to Repeated Burst protocol. **E**, fit of Spatiotemporal Discrete Release Model (*R*^2^ = 0.99) to Single Burst protocol. **F**, fit of Spatiotemporal Discrete Release Model (*R*^2^ = 0.96) to Repeated Burst protocol. Blue lines are model fits, blue triangles indicate start times *t*_*i*_ of electrical bursts, orange dots are averaged FSCV traces (*N* = 7 animals), and orange ribbons report SEM.

The long-term depression kinetic contributes to an ∼50% decrease in evoked DA release observed in WT mice between Sweeps 1 and 6. While the kick component of long-term depression is weaker than both short-term kinetics (Table 1), the time constant of 15 minutes is ∼60-fold longer than the closest time constant for short-term kinetics and consistent with the prior literature^9,17^. The release parameter estimates (Table 1) indicate that DAT activity *V*_*m*_ and DA release [*DA*]_*P*_ decrease by 33.3% and ∼25 – 30% between Sweeps 1 and 6. Further experiments can be conducted to investigate the biological mechanisms that relate to long-term plasticity of release and reuptake that could account for these effects, such as synaptic exhaustion at vesicular release sites and DAT endocytosis.

### Synuclein Proteins Regulate DA Release Kinetic Components In Vivo

In order to determine how the kinetic components can be altered in vivo, we analyzed the impact of the synuclein family of neuronal proteins on DA release. Synuclein proteins have three members: α-synuclein (α-Syn), β-synuclein (β-Syn), and γ-synuclein (γ-Syn)^18^. α-Syn and β-Syn are expressed in brain regions including the striatum, thalamus, hippocampus, neocortex, and cerebellum^19^, and γ-Syn is expressed in the peripheral nervous system as well as the brain^20^. α-Syn has been associated with disease-related protein aggregates in Parkinson’s disease and other neurodegenerative disorders, β-Syn has been shown to inhibit α-Syn aggregation, and abnormal levels of γ-Syn have been linked to multiple cancers^21^. Although the normal physiological functions of the synucleins are still under study, they regulate evoked DA release both in the striatal slice^22^ and in vivo^16^, likely by modulating steps in synaptic vesicle recycling and exocytosis^23,24^.

To analyze the interactions of synucleins on DA release in vivo and its effects on the kinetics of DA release, the models were fit to FSCV data from two lines of mutant mice: α-Syn knockout (KO) mice that are deficient in α-Syn, and synuclein triple knockout (TKO) mice that are deficient in α-Syn, β-Syn, and γ-Syn. Like WT mice, the data was taken from the Single Burst and Repeated Burst protocols for Sweep 1 and Sweep 6. Comparing the models’ kinetic parameter estimates (Table 1) between the WT mice (Figures 3 and 4), α-Syn KO mice (Figures 5 and 6), and Syn TKO mice (Figures 7 and 8) reveals that the kick component of short-term facilitation decreases by ∼40 – 50% and the time constant of short-term depression increases by ∼2.5-fold in the mutant mice, indicating that α-Syn expression decreases short-term depression and/or enhances short-term facilitation. In addition, the kick component of long-term depression is 0 in both α-Syn KO and Syn TKO mice, indicating that long-term depression is fully dependent on α-Syn expression. These results reinforce conclusions by Somayaji et al.^16^ that α-Syn controls the short-term facilitation and long-term depression of DA release.

**FIGURE 5:**
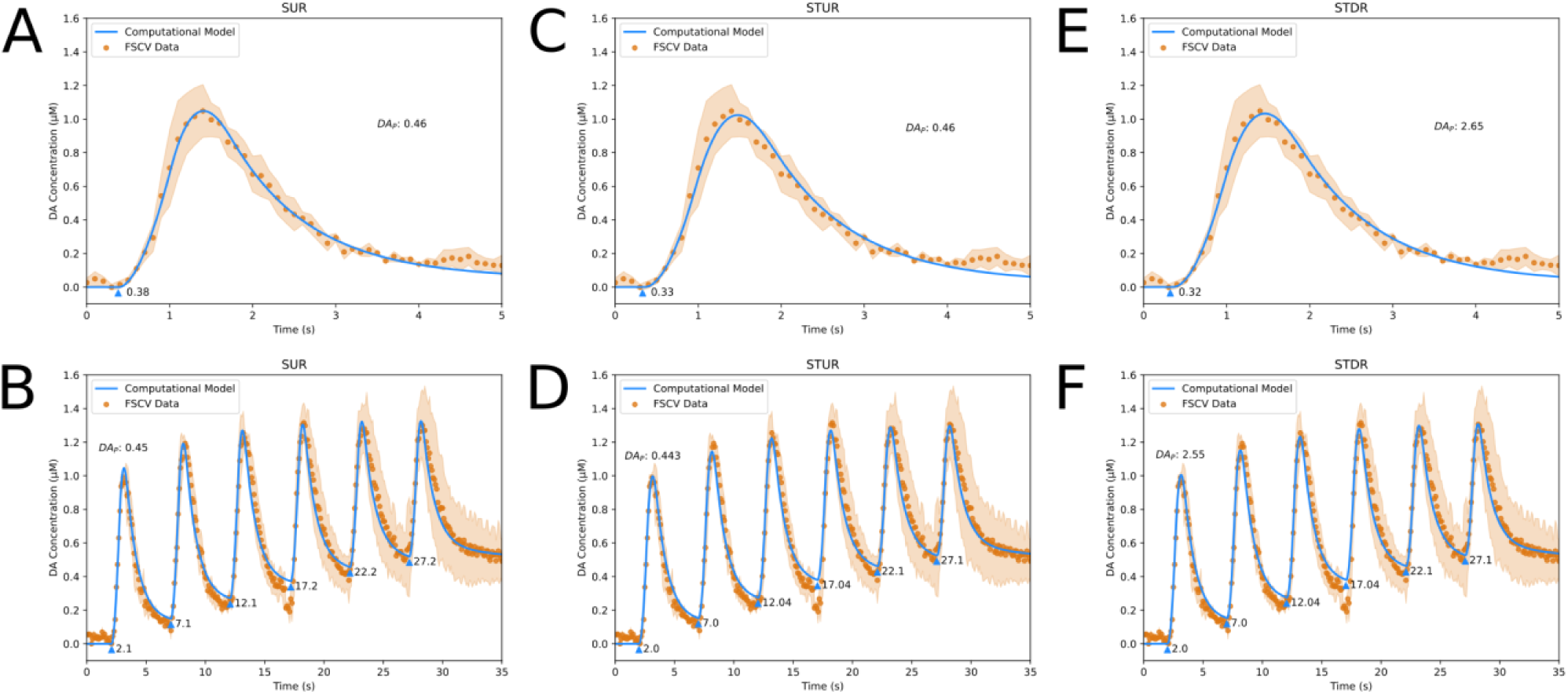
Fits of DA Computational Models to Sweep 1 FSCV Traces from α-Syn KO Mice in Dorsal Striatum. **A**, fit of Simple Uniform Release Model (*R*^2^ = 0.99) to Single Burst protocol. **B**, fit of Simple Uniform Release Model (*R*^2^ = 0.97) to Repeated Burst protocol. **C**, fit of Spatiotemporal Uniform Release Model (*R*^2^ = 0.98) to Single Burst protocol. **D**, fit of Spatiotemporal Uniform Release Model (*R*^2^ = 0.97) to Repeated Burst protocol. **E**, fit of Spatiotemporal Discrete Release Model (*R*^2^ = 0.98) to Single Burst protocol. **F**, fit of Spatiotemporal Discrete Release Model (*R*^2^ = 0.97) to Repeated Burst protocol. Blue lines are model fits, blue triangles indicate start times *t*_*i*_ of electrical bursts, orange dots are averaged FSCV traces (*N* = 3 animals), and orange ribbons report SEM.

**FIGURE 6:**
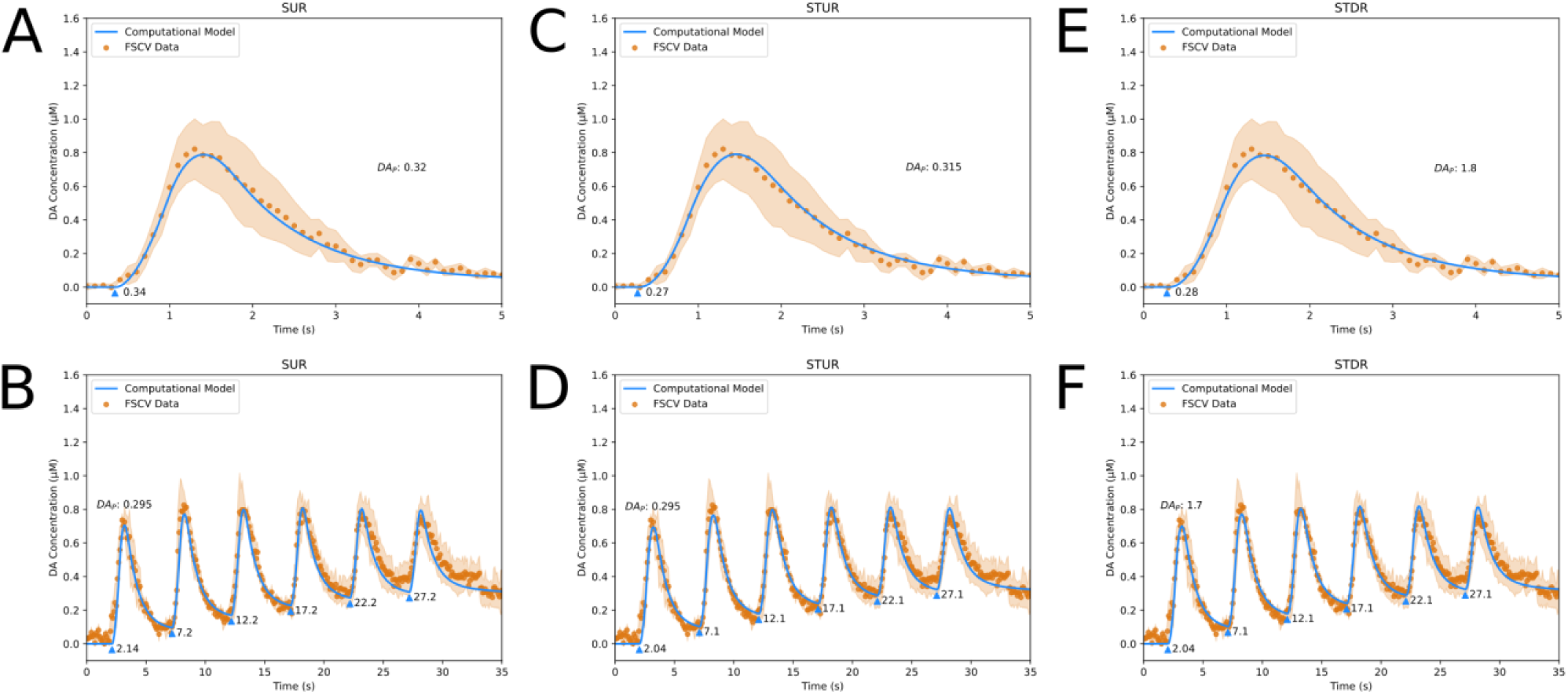
Fits of DA Computational Models to Sweep 6 FSCV Traces from α-Syn KO Mice in Dorsal Striatum. **A**, fit of Simple Uniform Release Model (*R*^2^ = 0.99) to Single Burst protocol. **B**, fit of Simple Uniform Release Model (*R*^2^ = 0.94) to Repeated Burst protocol. **C**, fit of Spatiotemporal Uniform Release Model (*R*^2^ = 0.99) to Single Burst protocol. **D**, fit of Spatiotemporal Uniform Release Model (*R*^2^ = 0.96) to Repeated Burst protocol. **E**, fit of Spatiotemporal Discrete Release Model (*R*^2^ = 0.99) to Single Burst protocol. **F**, fit of Spatiotemporal Discrete Release Model (*R*^2^ = 0.96) to Repeated Burst protocol. Blue lines are model fits, blue triangles indicate start times *t*_*i*_ of electrical bursts, orange dots are averaged FSCV traces (*N* = 3 animals), and orange ribbons report SEM.

**FIGURE 7:**
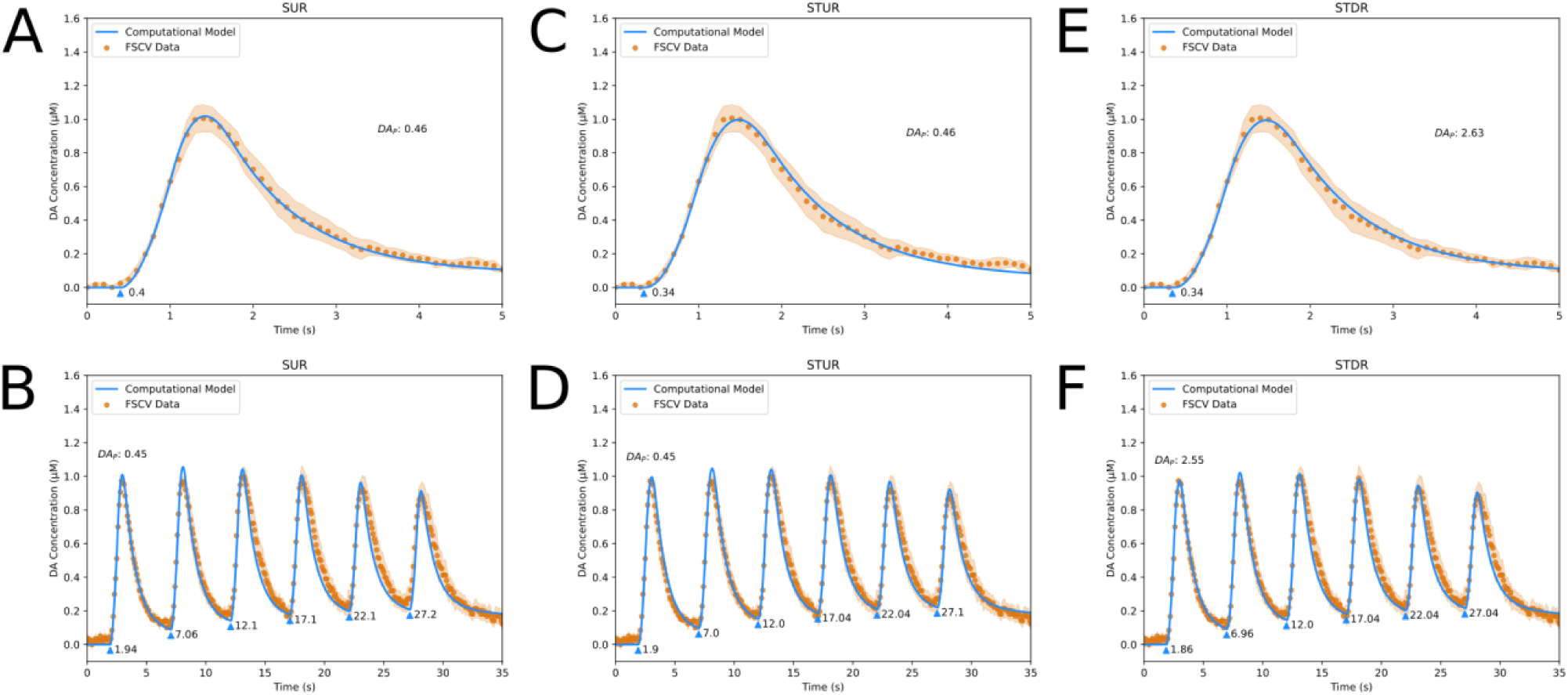
Fits of DA Computational Models to Sweep 1 FSCV Traces from Syn TKO Mice in Dorsal Striatum. **A**, fit of Simple Uniform Release Model (*R*^2^ = 1.00) to Single Burst protocol. **B**, fit of Simple Uniform Release Model (*R*^2^ = 0.95) to Repeated Burst protocol. **C**, fit of Spatiotemporal Uniform Release Model (*R*^2^ = 0.99) to Single Burst protocol. **D**, fit of Spatiotemporal Uniform Release Model (*R*^2^ = 0.97) to Repeated Burst protocol. **E**, fit of Spatiotemporal Discrete Release Model (*R*^2^ = 1.00) to Single Burst protocol. **F**, fit of Spatiotemporal Discrete Release Model (*R*^2^ = 0.97) to Repeated Burst protocol. Blue lines are model fits, blue triangles indicate start times *t*_*i*_ of electrical bursts, orange dots are averaged FSCV traces (*N* = 7 animals), and orange ribbons report SEM.

**FIGURE 8:**
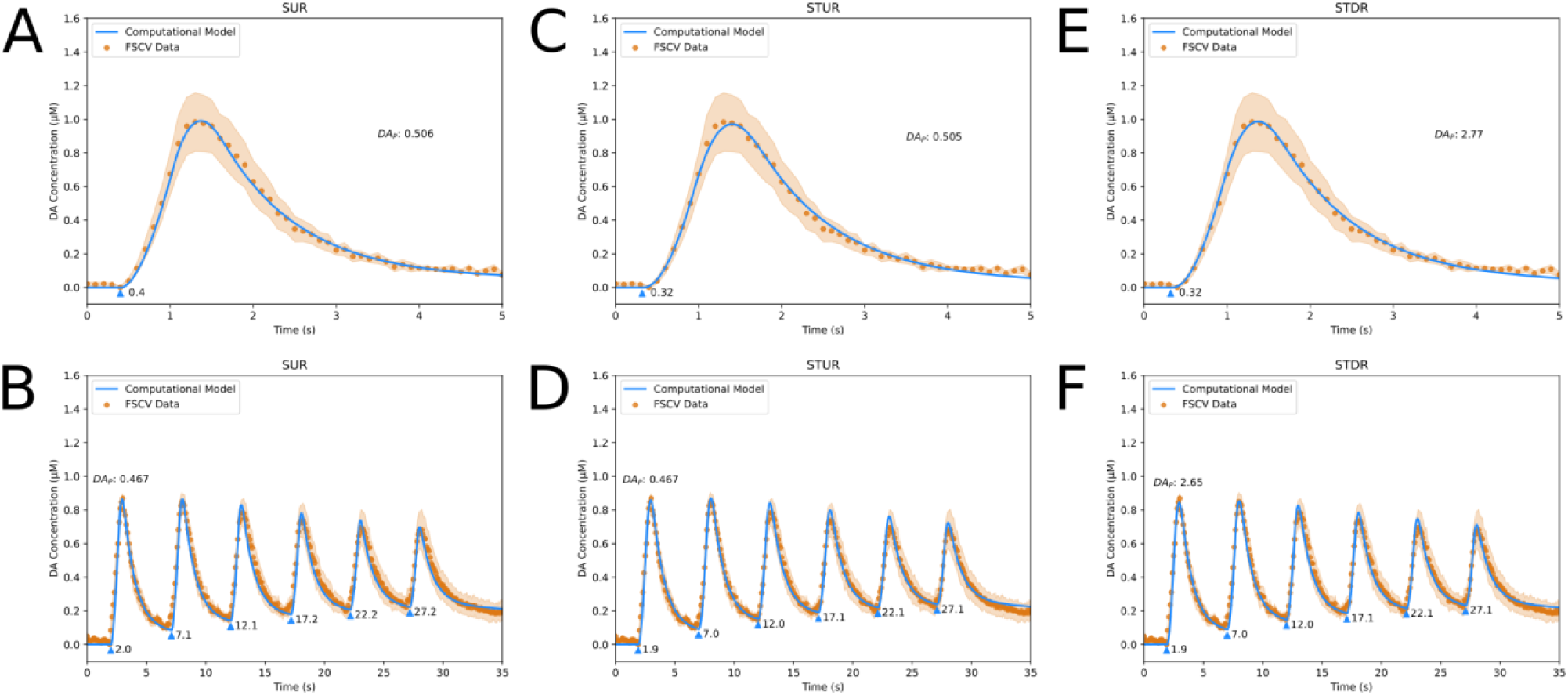
Fits of DA Computational Models to Sweep 6 FSCV Traces from Syn TKO Mice in Dorsal Striatum. **A**, fit of Simple Uniform Release Model (*R*^2^ = 1.00) to Single Burst protocol. **B**, fit of Simple Uniform Release Model (*R*^2^ = 0.96) to Repeated Burst protocol. **C**, fit of Spatiotemporal Uniform Release Model (*R*^2^ = 0.99) to Single Burst protocol. **D**, fit of Spatiotemporal Uniform Release Model (*R*^2^ = 0.98) to Repeated Burst protocol. **E**, fit of Spatiotemporal Discrete Release Model (*R*^2^ = 1.00) to Single Burst protocol. **F**, fit of Spatiotemporal Discrete Release Model (*R*^2^ = 0.98) to Repeated Burst protocol. Blue lines are model fits, blue triangles indicate start times *t*_*i*_ of electrical bursts, orange dots are averaged FSCV traces (*N* = 7 animals), and orange ribbons report SEM.

Similar to WT mice, the models’ release parameter estimates (Table 1) for α-Syn KO mice indicate a 33.3% decrease in DAT activity and ∼30 – 35% decrease in DA release between Sweeps 1 and 6, which accounts for the smaller ∼25% decrease in evoked DA release observed in the data between the sweeps in the absence of any long-term depression kinetic. In contrast, the models’ estimates for the Syn TKO mice indicate a relatively small *increase* in both DAT activity and DA release between Sweeps 1 and 6 by 16.7% and ∼5 – 10%, respectively (Table 1). This suggests a previously unknown role of β-Syn and/or γ-Syn in the long-term regulation of DAT uptake and vesicular DA release and may in part underlie the observation that the Syn TKO line has a greater effect than the α-Syn KO line on exocytosis of peptides from chromaffin cells and cultured hippocampal neurons^25^.

## Discussion

We presented three novel computational models of DA release to fit and analyze FSCV recordings of evoked DA release in the dorsal striatum of anesthetized mice. Using the models, we quantified biological mechanisms that influence DA release such as DAT uptake, DA diffusion, and DA release kinetics. As in a previous study of DA release during freely moving behavior^9^, we found that three kinetic components effectively describe modulation of DA release over time: short-term facilitation, short-term depression, and long-term depression. We also analyzed the complex interactions of the synuclein family of proteins with DA release using these models and mathematically quantified their effect. The results support our previous findings that α-Syn controls the short-term facilitation and long-term depression of DA. Furthermore, we identified a new role for β-Syn and/or γ-Syn in the long-term regulation of DAT uptake and DA release.

While FSCV experiments can be performed either in vivo or in the striatal slice, the computational models we present will not satisfactorily fit traces recorded in slice. The rapid recovery time of evoked DA release in vivo of < 10 milliseconds^16^ allows the electrical stimulation of midbrain neurons at 50 Hz, providing the means to study short-term synaptic plasticity. In contrast, the recovery time between electrical stimuli in slice is ∼1 minute^26^, which is due in part to the lack of tonic DA neuron firing and the lack of other neuronal activity in slice except for the cholinergic interneuron^27^. As such, single electrical pulses cannot be recorded in slice without an extended recovery time and rapid electrical pulse trains produce a negligible increase in DA release.

Given that the three computational models produce close fits to the data, they can be used without fundamentally altering the model’s estimates for the DA release kinetics or DAT uptake. For fitting data with multiple burst stimuli such as the Repeated Burst protocol, users may get slightly improved computational performance using the Simple Uniform Release Model. However, it may be beneficial to use the spatiotemporal models if the FSCV experiments are performed in conjunction with other electrochemical or optical procedures to elucidate physical characteristics in the striatum. For example, it is possible to use amperometry to estimate the size of the dead space around the electrode, and the current dead space radius of 3 μm specified for Figures 3 – 8 is derived from an amperometric study^10^. As the dead space will most often vary from experiment to experiment and can alter the DA measured by the carbon-fiber electrode (Figure 9), using amperometry and FSCV together can enable the dead space dimensions to be computed from amperometry data and used in the spatiotemporal models to fit FSCV data.

**FIGURE 9:**
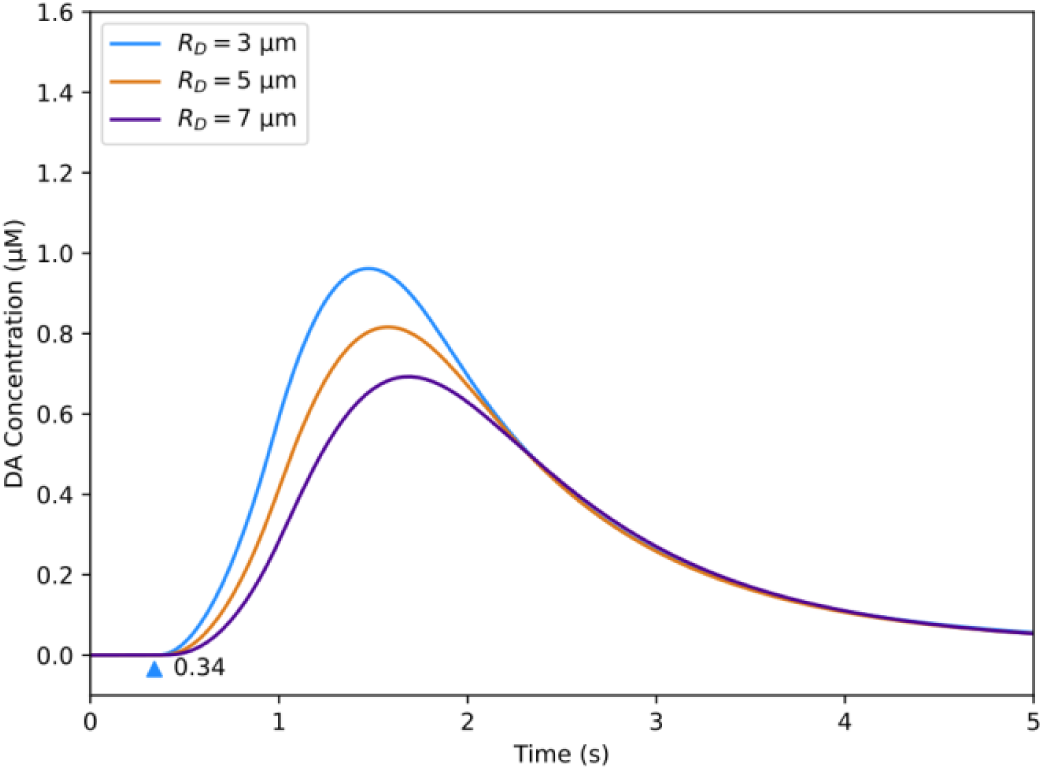
Effect of Dead Space Radius on Spatiotemporal Uniform Release Model Output. *R*_*D*_ is the parameter which controls the radius of the dead space in the model; 3 μm is the value used for fits presented in the Results section of the main paper. Blue triangle indicates the start time of the electrical burst. The remaining model parameters are set to values in Figure 3C (see Supplemental Table 1).

Additionally, non-electrochemical DA recording methods such as optical reporters or false fluorescent neurotransmitters (FFNs) can be used to directly visualize the distribution of active and silent release sites in the striatum, which would make the Spatiotemporal Discrete Release Model particularly beneficial compared to other models. Currently, the distance between two adjacent release sites in this model for the fits presented in Figures 3 – 8 is set to 6 μm based on numbers derived from false fluorescent neurotransmitter (FFN) imaging reported by Pereira et al^28^ but assumes that the release sites are spaced at equal intervals. In theory, if the FSCV experiments are performed in conjunction with FFN or optical reporter experiments, the position and distance of release sites can be adjusted to any location according to the experiment. While our analysis of FSCV provides genuine measures of DA concentration, in the future, it may be possible to adapt the models and describe kinetics of the change in fluorescence (ΔF/F) data from fluorescent reporters, as many labs have access to fiber photometry.

The cylindrical diffusion of DA implemented in the spatiotemporal models may be particularly useful for modeling DA release under the influence of DAT-inhibiting drugs like cocaine^29^ and nomifensine^30^. The diameter of the cylinder is set to 100 μm for the fits presented in Figures 3 – 8, less than the actual mouse striatum diameter of 2 – 3 mm^31^. Under typical circumstances, distant release sites do not contribute to the final FSCV trace because DAT will remove distally released DA before it diffuses to the electrode from the extracellular space. However, if DAT is at low levels or inhibited through pharmacological methods, the computed diameter of the striatum can be adjusted to capture DA diffusion in the extracellular space from distant release sites and account for the increase in DA concentration at the electrode.

## Methods

### Computational Models

The models were implemented in Python 3.8.5, and the detailed discretization schemes used in the implementation can be found in the Supplemental Information. All three models simulate the kinetics of DA release and the measurement of DA at the electrode using ordinary differential equations (ODEs).

The kinetics of stimulation-dependent DA release is implemented as a coupled ODE system revised from Montague et al.^9^ for anesthetized in vivo FSCV experiments. The ODEs simulate the kinetics as the product of “hidden” dynamic components *A*, and each component is a separate ODE *H*_*j*_.

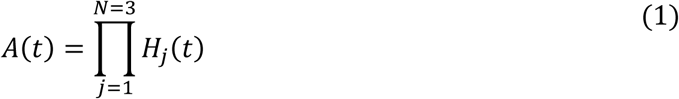

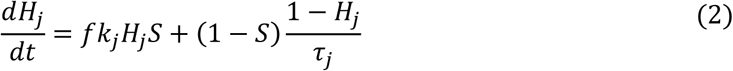

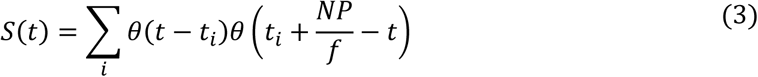

During stimulus periods, a kick component *K*_*j*_ is multiplied to each hidden component; positive values > 0 produce facilitation and negative values < 0 produce depression. During interstimulus periods, the hidden components decay towards the initial condition of 1 based on the time constant *τ*_*j*_. An example of the individual kinetic components is shown in Figure 10. The stimulation pattern *S* uses the Heaviside theta function *θ* to simulate electrical bursts, with *t*_*i*_ as the start time for each burst, *NP* as the number of pulses in each burst, and *f* as the stimulus frequency (in Hz). Due to the increased lag time introduced by spatial diffusion in the dead space, *t*_*i*_ is earlier in the STUR and STDR models compared to the SUR model.

**FIGURE 10:**
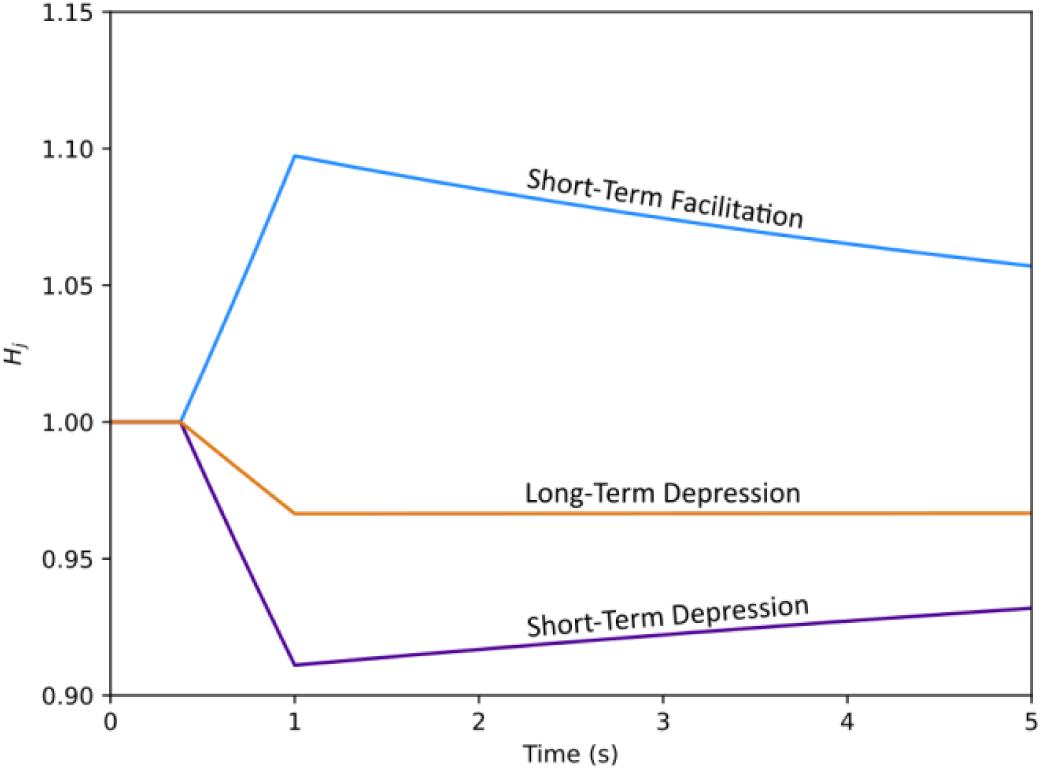
Example of Individual Kinetic Components (*H*_*j*_) in DA Computational Models. *H*_1_ (blue line) is short-term facilitation, *H*_2_ (purple) is short-term depression, and *H*_3_ (orange line) is long-term depression. The product of the three kinetic components determines the overall facilitative/depressive effect on the concentration of dopamine release. Refer to Equation 2 in the Methods section.

The measurement of DA at the electrode is also implemented as a coupled ODE system to model the slow temporal response and electrochemical adsorption that occur with the carbon-fiber electrode in FSCV experiments:

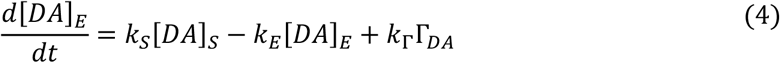

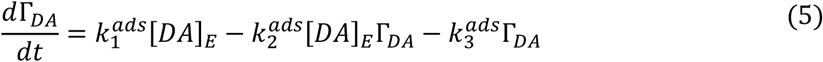

[*DA*]_*E*_ computes the DA diffusing from the extracellular space minus the DA leaving the electrode due to bounce back in FSCV experiments, plus the residual DA that occurs due to adsorption on the electrode. *K*_*S*_ is the rate transfer of DA moving from the striatum towards the electrode, *K*_*E*_ is the rate transfer of DA moving away from the electrode, and *K*_Γ_ = 1 is the rate transfer of DA adsorbing to the electrode. The adsorption Γ_*DA*_ is modeled as the concentration of DA that adsorbs to the electrode minus the concentration of DA that desorbs from the electrode, and it is a modified version of the original equation presented by Bath et al.^15^ The adsorption kinetic 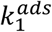 controls the amount of DA that adheres to the electrode, and the desorption kinetics 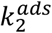 and 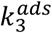 control the amount of residual DA on the electrode that falls off. The full derivation for Equation (5) is provided in the Supplementary Information.

### Simple Uniform Release Model

The DA released into the striatum [*DA*]_*S*_ is implemented as an ODE in the simple uniform release model.

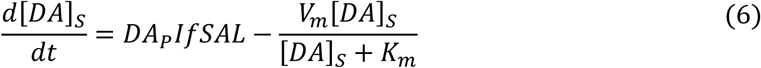

[*DA*]_*S*_ models the concentration of DA released by electrical stimulation into the striatum and removed through DAT uptake. *DA*_*P*_ is the amount of DA released per milliamp (mA), and *I* is the electrical stimulus current in mA, such that *DA*_*P*_*I* is the concentration of DA released per pulse. *V*_*m*_ and *K*_*m*_ are the maximal rate and affinity constant of DAT uptake^32^, modeled using first-order Michaelis-Menten kinetics^33^. A loss factor *L* < 1 is used to scale down the concentration of DA to account for the diffusion towards the electrode through the extracellular space and dead space.

### Spatiotemporal Uniform Release Model

A partial differential equation (PDE) is used to simulate spatiotemporal diffusion and assign a physical dimension to the dead space:

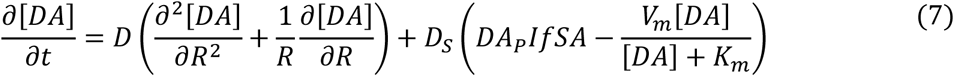

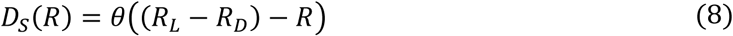

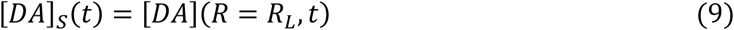

The striatum is modeled as a cylinder with uniformly distributed release sites, and the carbon-fiber electrode and dead space are positioned in the center of the striatum. The diffusion of DA in the striatum is simulated as isotropic diffusion using radial coordinates as described by John Crank^34^, where *R* signifies the distance along the radius of the cylinder and *D* = 240 μm^2^/s is the diffusion coefficient of DA in the brain accounting for tortuosity^35^. The dead space is computed as a new variable *D*_*S*_ using the Heaviside theta function *θ*, where *R*_*L*_ is the radius of the cylinder and *R*_*D*_ is the radius of the dead space. The area covered by the dead space does not include DA release and DAT uptake due to the damaged tissue^10^.

### Spatiotemporal Discrete Release Model

The release and diffusion of DA in this model are implemented nearly identically to the Spatiotemporal Uniform Release Model, except for the inclusion of an additional variable *P* to simulate spatially discrete release sites instead of a continuous release area:

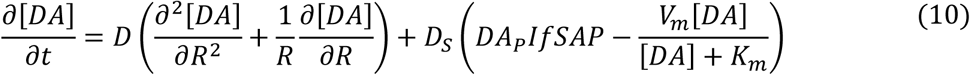

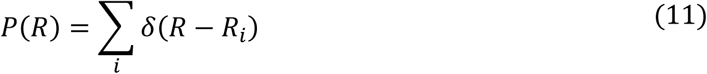

*P* uses the Dirac delta function *δ* to simulate multiple “point” sources of release in the cylinder, with *R*_*i*_ specifying the location of each point along the radius. Depending on the distribution of the release points, the estimates for *DA*_*P*_ can be higher compared to the Simple Uniform Release and Spatiotemporal Uniform Release models; sparser distributions lead to higher release from each individual point.

### Curve Fitting

Many of the current parameter estimation algorithms for ODE models are computationally intensive, and they typically necessitate the use of non-convex optimization approaches which require repeated numerical integration. Automatic differentiation variational inference (ADVI)^36,37^ is an efficient statistical inference algorithm that uses gradients to optimize the parameters and compute the best fit to the data. The algorithm works by transforming the parameters from the original latent variable space to the unconstrained latent variable space. It then uses both the unconstrained and original parameters to compute an objective called the evidence lower bound (ELBO), comprised of the expected log joint density, the log Jacobian of the transformation, and the entropy of the unconstrained parameters (which are drawn from a Gaussian distribution)^36^. The gradients of the unconstrained parameters with respect to the ELBO are used to optimize the original parameters, and the algorithm iterates until the parameters converge. Because the gradients can be computed efficiently using automatic differentiation, variational inference is typically faster than other statistical inference methods like Markov Chain Monte Carlo (MCMC)^38^.

Applying ADVI requires adapting the ODE model into a probabilistic model to compute the expected log joint density for the ELBO, comprised of the log-prior and log-likelihood. To calculate the log-prior, the free parameters in the ODE model 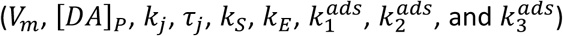 are drawn from independent Cauchy distributions. For the log-likelihood, the ODE’s solution at a given state 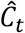 is computed by integrating [*DA*]_*E*_ over time, shown in Equation (12). 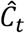 is then assigned as the mean of a Gaussian distribution to compute the log-likelihood (with the standard deviation *v* set to 0.05), and the corresponding state in the real data *C*_*t*_ is drawn from that distribution, shown in Equation (13):

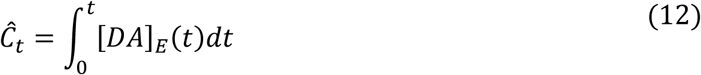

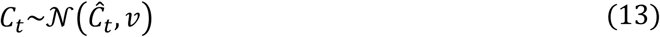

### In Vivo Recordings

To validate the computational models in this article, the in vivo FSCV traces of DA release and the detailed experimental procedures used were previously published^16^. Briefly, male or female mice were anesthetized with isoflurane (induction 2.5%, maintenance 0.8 – 1.4% in O_2_, 0.35 L/min), placed on a heating pad, and head-fixed on a stereotaxic frame. A craniotomy was performed to insert a 22G bipolar stimulating electrode (P1 Technologies) in the ventral midbrain to trigger electrical pulse trains and a carbon fiber microelectrode (5 μm diameter, ∼150 μm length) in the dorsal striatum to measure evoked DA release; detailed stereotaxic coordinates and procedures can be found in the publication. The depth of the stimulating electrode was adjusted between 4 – 4.5 mm for maximal DA release. To conduct FSCV, a triangular voltage waveform (–450 to +800 mV at 294 mV/ms versus Ag/AgCl reference electrode) was applied to the carbon fiber electrode at a 10 Hz sampling rate (or every 100 ms). The signals measured at the electrode were converted from analog to digital signals using a 16-bit data acquisition interface and subsequently recorded using IGOR Pro-6.37 software. The electrical pulse trains were delivered to the stimulating electrode at a constant current of 400 μA using a stimulus isolator and a pulse generator. The stimulation protocols used in the experiments are explained in the Results section. The carbon fiber microelectrodes were calibrated in artificial cerebrospinal fluid using known concentrations of DA.

## Acknowledgments

We would like to thank Dr. Mark Wightman for his inputs on the computational models and Dr. Charles Nicholson for his inputs on the computation of the dead space and diffusional geometry and comments on the manuscript. This research was supported in part by the NSF GRFP grant no. DGE 2036197 (SN), European Research Council Synergy grant ASTRA no. 855923 (MM and GR), NIH grants R01DA007418 and R01MH108186 (DLS), and the JPB foundation (DLS).

## Author Contributions

SN and DLS designed the research. SN, MM, and GR designed computational models with input from EVM and DLS. MS performed in vivo FSCV experiments. SN and MM implemented computational models. SN and DAK designed curve fitting algorithm. SN implemented curve fitting algorithm and performed model fitting analysis. SN and DLS wrote the manuscript, with all authors providing input.

