## Supplemental Information for "Novel Computational Models of Evoked Dopamine Release In Vivo Measured by Fast Scan Cyclic Voltammetry Quantify the Regulation of Presynaptic Kinetics by Synucleins"

### Model Implementation

The differential equations in the models were integrated using finite-difference discretization schemes. A finite-difference scheme discretizes a function into a grid by partitioning the temporal and spatial dimensions into evenly-spaced time points and space points, and the derivatives are computed at each time and space point to approximate the solution. In the Simple Uniform Release Model which is only integrated over time, the time step interval $\Delta t = \frac{1}{f}$, with $f$ as the stimulation frequency in the FSCV experiments (50 Hz in the current data). In the spatiotemporal models, the time step $\Delta t = \frac{(\Delta R)^2}{2D}$ is computed based on the space step and the diffusion coefficient[1], where the space step $\Delta R$ is set to 1 µm and the diffusion coefficient for DA $D$ is set to 240 µm$^2$/s based on the literature[2].

The ordinary differential equations (ODEs) in the computational models were discretized using the forward Euler scheme:

$$H_j(t+\Delta t) = H_j(t) + \Delta t \left[ f k_j H_j(t) S(t) + \{1 - S(t)\} \frac{1 - H_j(t)}{\tau_j} \right] \tag{1}$$

$$[DA]_E(t+\Delta t) = [DA]_E(t) + \Delta t [k_S [DA]_S(t) - k_E [DA]_E(t) + k_\Gamma \Gamma_{DA}(t)] \tag{2}$$

$$\Gamma_{DA}(t+\Delta t) = \Gamma_{DA}(t) + \Delta t \left[ k_1^{ads} [DA]_E(t) - k_2^{ads} [DA]_E(t) \Gamma_{DA}(t) - k_3^{ads} \Gamma_{DA}(t) \right] \tag{3}$$

$$[DA]_S(t+\Delta t) = [DA]_S(t) + \Delta t \left[ DA_P I f S(t) A(t) L - \frac{V_m [DA]_S(t)}{[DA]_S(t) + K_m} \right] \tag{4}$$

For the partial differential equation (PDE) in the Spatiotemporal Uniform Release Model (Equation (5)) and the Spatiotemporal Discrete Release Model (Equation (6)), the cylindrical diffusion discretization was adapted from Venton et al.[1] using an explicit finite-difference method, and the DA release and reuptake were adapted from the ODE discretization schemes:

$$[DA](t+\Delta t, R) = [DA](t,R) + C_D + D_s(R)[DA_P I f S(t) A(t) - M]\Delta t \tag{5}$$

$$[DA](t+\Delta t, R) = [DA](t,R) + C_D + D_s(R)[DA_P I f S(t) A(t) P(R) - M]\Delta t \tag{6}$$

$$C_D = D \left[ \frac{R}{2R-1} [DA](t, R+\Delta R) + \frac{R-1}{2R-1} [DA](t, R-\Delta R) - [DA](t,R) \right] \Delta t \tag{7}$$

$$M = \frac{V_m [DA](t,R)}{[DA](t,R) + K_m} \tag{8}$$

**DA Adsorption**

The concentration of DA that occurs due to electrochemical adsorption $\Gamma_{[DA]}$ is derived from an ODE that describes the number of molecules that are adsorbed by the electrode over time:

$$\frac{dN_{ads}}{dt} = [DA]_E \frac{l}{t_1}(\Sigma_0 - \sigma N_{ads}) - \frac{N_{ads}}{t_2} \tag{9}$$

$N_{ads}$ is the net difference between the number of molecules that adsorb and desorb to the electrode. $[DA]_E$ describes the concentration of DA near the electrode (see Equation 4 in the Methods section). $l$ is the width of the layer of molecules that are in contact with the electrode, $\Sigma_0$ is the total surface area of the electrode, and $\sigma$ is the surface area occupied by one molecule, such that $\Sigma_0 - \sigma N_{ads}$ is the free surface of the electrode. $t_1$ and $t_2$ are time constants that control the adsorption and desorption time, respectively. To recast this equation in terms of concentration, all of the terms can be divided by $V = Bl$, with $B$ having the dimensions of a surface:

$$\frac{d\Gamma_{[DA]}}{dt} = k_1^{ads}[DA]_E - k_2^{ads}[DA]_E\Gamma_{[DA]} - k_3^{ads}\Gamma_{[DA]} \tag{10}$$

where $\Gamma_{[DA]} = \frac{N_{ads}}{lB}$, $k_1^{ads} = \frac{\Sigma_0}{Bt_1}$, $k_2^{ads} = \frac{\sigma l}{t_1}$, and $k_3^{ads} = \frac{1}{t_2}$.

| | $DA_P$ (µM/mA) | $V_m$ (µM/s) | $K_m$ (µM) | $k_S$ ($s^{-1}$) | $k_E$ ($s^{-1}$) | $k_1^{ads}$ ($s^{-1}$) | $k_2^{ads}$ ($s^{-1}$) | $k_3^{ads}$ ($s^{-1}$) |
|---|---|---|---|---|---|---|---|---|
| **Fig 3A** | 0.420 | 4.8 | 0.2 | 0.9 | 1.05 | 0.035 | 0.140 | 0.090 |
| **Fig 3B** | 0.395 | 4.8 | 0.2 | 0.9 | 0.90 | 0.035 | 0.140 | 0.090 |
| **Fig 4A** | 0.305 | 3.2 | 0.2 | 0.9 | 1.00 | 0.035 | 0.140 | 0.090 |
| **Fig 4B** | 0.280 | 3.2 | 0.2 | 0.9 | 0.90 | 0.045 | 0.140 | 0.090 |
| **Fig 5A** | 0.460 | 4.8 | 0.2 | 0.9 | 1.00 | 0.040 | 0.140 | 0.090 |
| **Fig 5B** | 0.450 | 4.8 | 0.2 | 0.9 | 0.90 | 0.070 | 0.037 | 0.070 |
| **Fig 6A** | 0.320 | 3.2 | 0.2 | 0.9 | 1.00 | 0.035 | 0.140 | 0.090 |
| **Fig 6B** | 0.295 | 3.2 | 0.2 | 0.9 | 0.90 | 0.065 | 0.050 | 0.070 |
| **Fig 7A** | 0.460 | 4.8 | 0.2 | 0.9 | 1.00 | 0.055 | 0.020 | 0.075 |
| **Fig 7B** | 0.450 | 4.8 | 0.2 | 0.9 | 0.90 | 0.040 | 0.050 | 0.070 |
| **Fig 8A** | 0.506 | 5.6 | 0.2 | 0.9 | 1.00 | 0.035 | 0.020 | 0.075 |
| **Fig 8B** | 0.467 | 5.6 | 0.2 | 0.9 | 0.90 | 0.050 | 0.020 | 0.075 |

**SUPPLEMENTAL TABLE 1:** Best Fit Parameters of Simple Uniform Release Model. Across all figures, $I$ = 0.4 mA, $f$ = 50 Hz, $NP$ = 30 pulses, and $L$ = 0.9.

| | $DA_P$ (μM/mA) | $V_m$ (μM/s) | $K_m$ (μM) | $k_S$ (s$^{-1}$) | $k_E$ (s$^{-1}$) | $k_1^{ads}$ (s$^{-1}$) | $k_2^{ads}$ (s$^{-1}$) | $k_3^{ads}$ (s$^{-1}$) |
|---|---|---|---|---|---|---|---|---|
| **Fig 3C** | 0.422 | 4.8 | 0.2 | 0.9 | 1.05 | 0.025 | 0.140 | 0.090 |
| **Fig 3D** | 0.400 | 4.8 | 0.2 | 0.9 | 0.90 | 0.035 | 0.140 | 0.090 |
| **Fig 4C** | 0.310 | 3.2 | 0.2 | 0.9 | 1.00 | 0.035 | 0.140 | 0.090 |
| **Fig 4D** | 0.290 | 3.2 | 0.2 | 0.9 | 0.90 | 0.045 | 0.140 | 0.090 |
| **Fig 5C** | 0.460 | 4.8 | 0.2 | 0.9 | 1.00 | 0.020 | 0.140 | 0.090 |
| **Fig 5D** | 0.443 | 4.8 | 0.2 | 0.9 | 0.90 | 0.070 | 0.037 | 0.070 |
| **Fig 6C** | 0.315 | 3.2 | 0.2 | 0.9 | 1.00 | 0.035 | 0.140 | 0.090 |
| **Fig 6D** | 0.295 | 3.2 | 0.2 | 0.9 | 0.90 | 0.065 | 0.050 | 0.070 |
| **Fig 7C** | 0.460 | 4.8 | 0.2 | 0.9 | 1.00 | 0.035 | 0.020 | 0.075 |
| **Fig 7D** | 0.450 | 4.8 | 0.2 | 0.9 | 0.90 | 0.040 | 0.050 | 0.070 |
| **Fig 8C** | 0.505 | 5.6 | 0.2 | 0.9 | 1.00 | 0.020 | 0.020 | 0.075 |
| **Fig 8D** | 0.467 | 5.6 | 0.2 | 0.9 | 0.90 | 0.050 | 0.020 | 0.075 |

**SUPPLEMENTAL TABLE 2:** Best Fit Parameters of Spatiotemporal Uniform Release Model. Across all figures, $I$ = 0.4 mA, $f$ = 50 Hz, $NP$ = 30 pulses, $R_L$ = 50 μm, and $R_D$ = 3 μm.

| | $DA_P$ (μM×μm/mA) | $V_m$ (μM/s) | $K_m$ (μM) | $k_S$ ($s^{-1}$) | $k_E$ ($s^{-1}$) | $k_1^{ads}$ ($s^{-1}$) | $k_2^{ads}$ ($s^{-1}$) | $k_3^{ads}$ ($s^{-1}$) |
|---|---|---|---|---|---|---|---|---|
| **Fig 3E** | 2.43 | 4.8 | 0.2 | 0.9 | 1.05 | 0.025 | 0.140 | 0.090 |
| **Fig 3F** | 2.30 | 4.8 | 0.2 | 0.9 | 0.90 | 0.035 | 0.140 | 0.090 |
| **Fig 4E** | 1.79 | 3.2 | 0.2 | 0.9 | 1.00 | 0.035 | 0.140 | 0.090 |
| **Fig 4F** | 1.70 | 3.2 | 0.2 | 0.9 | 0.90 | 0.045 | 0.140 | 0.090 |
| **Fig 5E** | 2.65 | 4.8 | 0.2 | 0.9 | 1.00 | 0.020 | 0.140 | 0.090 |
| **Fig 5F** | 2.55 | 4.8 | 0.2 | 0.9 | 0.90 | 0.070 | 0.037 | 0.070 |
| **Fig 6E** | 1.80 | 3.2 | 0.2 | 0.9 | 1.00 | 0.035 | 0.140 | 0.090 |
| **Fig 6F** | 1.70 | 3.2 | 0.2 | 0.9 | 0.90 | 0.065 | 0.050 | 0.070 |
| **Fig 7E** | 2.63 | 4.8 | 0.2 | 0.9 | 1.00 | 0.055 | 0.020 | 0.075 |
| **Fig 7F** | 2.55 | 4.8 | 0.2 | 0.9 | 0.90 | 0.040 | 0.050 | 0.070 |
| **Fig 8E** | 2.77 | 5.6 | 0.2 | 1.0 | 1.00 | 0.020 | 0.020 | 0.075 |
| **Fig 8F** | 2.65 | 5.6 | 0.2 | 0.9 | 0.90 | 0.050 | 0.020 | 0.075 |

**SUPPLEMENTAL TABLE 3:** Best Fit Parameters of Spatiotemporal Discrete Release Model. Across all figures, $I$ = 0.4 mA, $f$ = 50 Hz, $NP$ = 30 pulses, $R_L$ = 50 μm, and $R_D$ = 3 μm.
